# Resource availability and predator cues shape within- and transgenerational reaction norms

**DOI:** 10.64898/2025.12.05.692509

**Authors:** Léo Dejeux, Juliette Tariel-Adam, Sandrine Plénet, Emilien Luquet

## Abstract

The main dilemma facing prey is how to reduce predation risk while acquiring resources. Phenotypic plasticity, by modulating traits for acquiring resources and for defences, is a key process driving this trade-off. While there is ample evidence that predator-induced within-generational plasticity depends on resources, the relationship between resources and predator-induced transgenerational plasticity remains understudied. This study investigated how predator exposure and resource availability influence within- and trans-generational plasticity. We based our predictions on theories related to information- and state-based plasticity, two kinds of plasticity occurring in response to environmental information or alteration affecting somatic state, respectively. We conducted a two-generation laboratory experiment in the freshwater snail *Physa acuta*, manipulating predator cues and resource availability. Our results revealed that both types of plasticity participate in shaping the within- and transgenerational responses. Within a generation, the limited resource decreased the somatic state but did not constrain the defences. Across generations, the poor parental state decreased the offspring state even in high resource, which in turn decreased the absolute levels of their defences but increased their investment in reproduction. These results highlight the importance to integrate informational and state-based effects across generations to provide a more complete understanding of prey responses to predators.

## Introduction

Phenotypic plasticity, the capacity of organisms to produce different phenotypes depending on environmental conditions [1], is at the center of the most fundamental trade-offs prey face: the dilemma between acquiring resources or ensuring safety (e.g. [2–5]). Prey can perceive predator cues (mechanical, visual, auditory, chemical) and plastically develop anti-predator traits (behaviour, morphology, life history; [6]). The availability of resources in the environment influences resource acquisition by shaping how prey plastically adjust their phenotype to forage and ultimately determines their somatic state [7]. In environments combining risk of predation and limited resources, the prey phenotype results from a trade-off between traits beneficial for defending prey against predators or acquiring resources. For example, the trade-off in many organisms between foraging behavior and anti-predator morphology (review in [8]), the trade-off in *Rana sylvatica* tadpoles between growth (driven by gut size) and anti-predator morphology (tail size; [9]) or yet the trade-off in *Helisoma trivolis* freshwater snails between growth and anti-predator behaviour [10], are all influenced by resource availability in the environment.

While for a long time, phenotypic plasticity has only been considered within the lifetime of the organisms (within-generational plasticity, WGP), more recently, plasticity is widely recognized to occur across generations with the phenotype of a generation influenced by the environment experienced by the previous generation(s) regardless of underlying mechanisms (i.e. the broad-sense definition of transgenerational plasticity, TGP; [11,12]). Given the importance of resource availability as a mediator of predator-induced WGP, one would expect it to be important to TGP. However, the relationship between resource availability and predator-induced TGP is understudied. Recent syntheses on predator-induced TGP do not discuss resource manipulations or seem to include any studies that had resource manipulation[13,14]. Our study aims to fill this gap, investigating how the combination of both predator exposure and resource availability influence WGP and TGP through the resource acquisition-safety trade-off.

Predator exposure and resource availability have often been associated with two types of plastic responses: information-based and state-based plasticity, respectively [15]. Information-based plasticity is triggered by the detection of environmental cues informing individuals about the change in selective environmental conditions (e.g. predator cues). Then, individuals shape their own phenotype in accordance (e.g. anti-predator traits) and/or pass on the information to their offspring (e.g. presence of predators) regardless of their own parental somatic state, thereby influencing offspring phenotype (e.g. anti-predator traits) to better anticipate the likely future environmental conditions (e.g. predator-risk environment; that’s why information-based TGP is also called ‘anticipatory parental effect’; [11,16–18]). In that case, it is predicted that fitness is the highest when the early/parental and late/offspring environments match (i.e. when cues reliably predict the selective pressure, [15,19–21]), resulting in non-additive patterns between the effects of early/parental and late/offspring environments (environmental matching; [16]).

State-based plasticity reflects that environment (e.g. resource availability) may influence the somatic state of individuals regardless of the detection of reliable environmental information [18,22–24]. Although traditionally perceived as non-adaptive responses (i.e. physiological constraints), it is now widely recognized that state-based plasticity can also be adaptive, improving parental and/or offspring fitness, with individuals adjusting their performance and that of their offspring based on the quality of the environment they have experienced (that’s why state-based TGP is also called ‘parental condition-transfer effect’; [22,23,25,26]). In this type of plasticity, one can expected that good-state organisms outperform poor-state ones and produce good-state offspring that outperform offspring from poor-state parents regardless of the environment that offspring experience [23].

In general, disentangling between information- and state-based plasticity is challenging because they can operate at the same time, leading to non-additive patterns [18,22]. This is especially true in the context of resource acquisition-safety trade-off as defensive traits are costly to produce (e.g. [27]). The individual’s state can determine the amount of energy to allocate towards the response to predator cues [28] or conversely the response to predator cues can influence individual’s state (e.g. as a consequence of stress or changes in foraging; e.g. [2]). In addition, in some examples, the change in the somatic state may be an anti-predator trait, intertwining information-based and state-based plasticity. For example, prey can increase growth to reach a refuge size limiting predator attacks [29,30]. In this study, rather than attempting to disentangle information- and state-based plasticity, we merged the theoretical predictions of information- and state-based plasticity to predict how the combination of both predator exposure and resource availability influence WGP and TGP. <u>We</u> proposed three theoretical patterns of reaction norms (Fig. 1). These patterns assume that (i) the individuals developing in a low-quality environment (e.g. low resource availability) will be in a poorer somatic state and (ii) the anti-predator traits are costly to produce (i.e. condition-dependent expression; [31–33]). A first pattern is the absence of interaction between information- and state-based plasticity (‘additive pattern’; Fig. 1a): the anti-predator traits are induced by current or parental predator cues independently of the somatic state. Second, the state-based plasticity may constrain the information-based plasticity by reducing the slope of the reaction norm in poor environments (‘masking pattern’; Fig. 1b). The poor somatic state of individuals may for example limit the energy allocation towards the induction of anti-predator traits. Third, state-based plasticity may reveal the information-based responses by increasing the slope of the reaction norm in poor environments (‘revealing pattern’; Fig. 1c). Despite a poor somatic state in a poor environment, individuals still induce anti-predator traits. The opposite ‘masking’ and ‘revealing’ patterns may result from a trade-off between growth, reproduction and anti-predator traits; a trade-off that may give the best fitness outcome given the environmental conditions and hence be favoured by selection.

**Figure 1.**
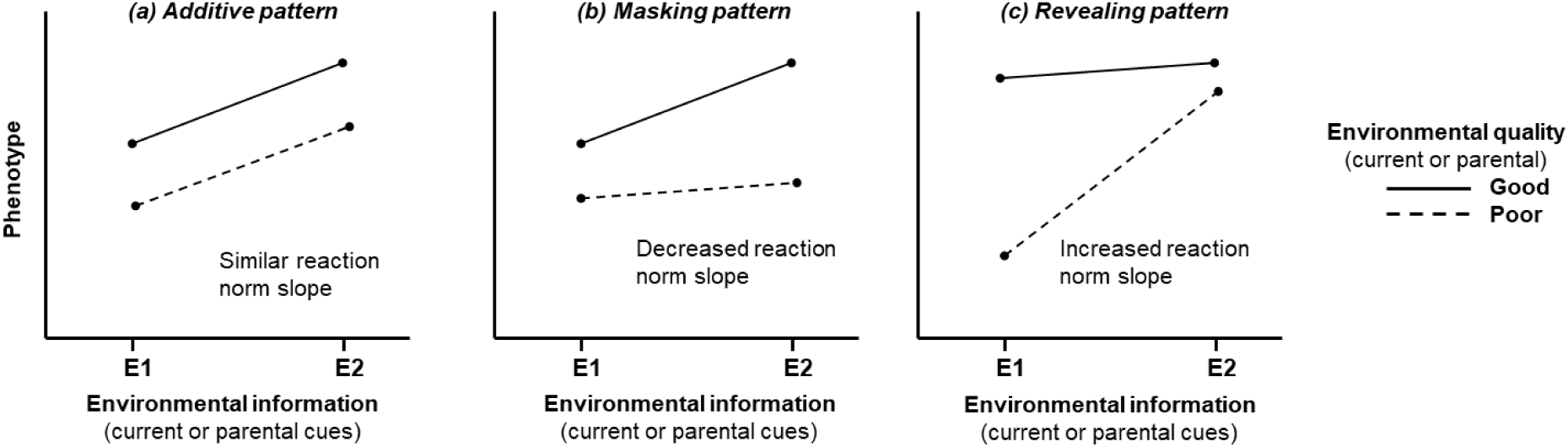
Theoretical patterns of reaction norms in the context of an interaction between information- and state-based plasticity (WGP or TGP). a) Additive pattern: absence of interaction between the effects of environmental information and quality. b) Masking pattern: the environmental quality may constrain the somatic state and consequently the plastic response to environmental information by reducing the slope of the reaction norm. c) Revealing pattern: the environmental quality may reveal the plastic response to environmental information by increasing the slope of the reaction norm. The y-axis shows the phenotype of the individuals (WGP) or of their offspring (TGP) according to the environmental information (x-axis: E1, E2) and quality (Poor or Good, dashed or full line respectively) for the current or the parental generation respectively.

We used the freshwater snail *Physa acuta* – crayfish *Faxonius limosus* system, a well-documented system of predator-induced WGP and TGP (e.g. [34–36]). We investigated how exposure to low resource availability and predator cues influenced the WGP reaction norm of exposed individuals and the TGP reaction norm of their offspring reared in a favourable environment (*ad libitum* resources and no predator exposure). We measured growth traits (shell size and total mass) through development (juvenile and adult stages), anti-predator traits (escape behaviour, shell-crush resistance; [37–39]) and reproductive outputs (number of eggs, number of juveniles). We expected that information- and state-based plasticity will interact both within and across generations according to a masking or revealing scenario (Fig. 1b, c).

## Materials and Methods

### Snail origin

*Physa acuta* (Physidae, Hygrophila, Gastropoda) is a freshwater snail with a worldwide geographical distribution [40]. It is a simultaneous hermaphrodite with preferential outcrossing, *i.e.* eggs are self-fertilized only when no mate is available to provide allosperm [41]. In this study, adult *P. acuta* snails from a laboratory outbred population have been used as source (F0 individuals). These snails originated from wild populations from lentic backwater of the Rhône River near Lyon (southeastern France) and have been reared in controlled conditions (25°C and a 12-hour light/dark cycle) for 21 generations. 112 F0 adult individuals were grouped together for three days in two 10 L aquariums filled with dechlorinated tap water for mating and laid eggs before being removed. Such mass mixed breeding approach reflects the breeding in the natural environment (i.e. repeated breeding as the opportunity arise) but prevent the recording of family identity. The eggs hatched after approximately seven days at 25°C and constituted the F1 generation.

### Experimental design

#### F1 generation

F1 individuals developed during 10 days in dechlorinated tap water in the 10 L aquariums. Next, 112 F1 snails were randomly assigned to each of the four treatments: control water (i.e. without predator-cues) with *ad libitum* food (Control non-exposed and High resource, CH), control water with restricted food (Control non-exposed and Low resource, CL), predator-cue exposure with *ad libitum* food (Predator-cues and High resource, PH) and predator-cue exposure with restricted food (Predator-cues and Low resource, PL), resulting in a full-factorial design (n total = 112 snails * 2 predator-cues conditions * 2 resource conditions = 448 snails). Each individual was isolated in an 80 mL box, the water was renewed twice a week, and snails were fed with chopped boiled lettuce. Exposure to predator cues was achieved by rearing snails in water containing predator cues. This predator-cue water was obtained by housing *Faxonius limosus* crayfish (one crayfish per 5 L) that were fed with snails for three days. After collecting the crayfish’s water, we added ca. 1.5 g of live *P. acuta* snails crushed for 1h in the solution. Thus, the predator-cue water contained olfactory cues of crayfish presence, both crayfish kairomones and snail alarm cues [42]. While for high-resource treatment, snails were fed *ad libitum* at each water renewal (twice a week), low-resource treatment was achieved by feeding the snails only once a week during the first water renewal (no food was added at the second water renewal, leaving the snails without food for the next 3 days). This alternating availability and unavailability of resources is consistent with natural conditions (biofilm availability varies in space and times in freshwater ecosystems) and allows for precise control over resource availability whilst simulating low resource availability throughout the snails’ lifetimes. At 25 days (15 days of treatment, called ‘juvenile stage’ thereafter), total mass and shell length were measured for each individual and then snails were put back in their boxes to continue the experiment. At 45 days old (35 days of treatment, called ‘adult stage’ thereafter), crawling-out of the water behaviour was recorded, which is a classic response to benthic predators such as crayfish [42,43]. Snails were again measured (total mass, shell length) and grouped together in 10 L aquarium within each treatment to mate for 24 hours (i.e. mass mating allowing random copulations). Then, F1 snails were isolated to lay eggs for 3 days, allowing us to assess the number of eggs laid and the number of juveniles produced, which subsequently became the F2 generation. Finally, F1 snails were sacrificed - put in boiling water to extract soft tissues of the shell - to subsequent shell-crush resistance measure, a defence against crayfish predation [34,35,39].

#### F2 generation

The F2 experimental design was the same than for F1, except that F2 snails were all reared in control conditions (control non-exposed and high resource, i.e. in a water without predator-cues and with *ad libitum* food): at 10 days old, F2 individuals within each parental treatment were pooled in aquarium, then 112 F2 snails per parental treatment were randomly selected and isolated (n total = 112 F2 snails * 2 parental predator-cues conditions * 2 parental resource conditions = 448 F2 snails) and finally measured at the ‘juvenile stage’ (i.e. 25 days; total mass and shell length) and at the ‘adult stage’ (i.e. 45 days; crawling-out behaviour, total mass, shell length, shell crush resistance, number of eggs laid and number of juveniles produced).

The experiment was conducted in a temperature-controlled room, maintaining a constant temperature of 25°C and a 12-hour light/dark cycle.

### Trait measurements

***Snail mass at juvenile and adult ages***: the total fresh mass of each snail (shell mass + body mass) was determined using a precision scale the nearest 0.001 g. Prior to measurement, snails were gently dried with tissue to remove excess moisture. ***Shell length at juvenile and adult stages***: the maximum length of the snail shell was measured using ImageJ software [44] at the nearest 0.001 mm from a photograph of the shell with its aperture upward taken by a camera mounted on a binocular loop. ***Crawling-out behaviour*** was scored once at the adult stage (45 days old) as a binary variable (the snail was out of the water or touching the surface versus in the water; [39]), approximately three hours after the water renewal. ***Shell crush resistance at adult age***: the resistance of empty shells to crushing was assessed by a device designed to apply pressure on the shell (with aperture pointing downward) and recorded the force at the nearest 0.1 g. The force recorded at the peak represents the force needed to crush the shell (see [39] for more details). ***The number of eggs*** laid by each snail were counted before hatching. ***The number of juveniles*** was counted 10 days after laying (i.e. approx. 2-3 days after hatching).

### Statistical analyses

Statistical analyses tested the effects of predator-exposure, resource availability and their interaction on the expression of traits in parents (F1) and in offspring (F2). Data from the F1 and F2 generations were analysed separately. ***Snail size:*** shell length and snail mass (cube root transformed) were highly correlated in both F1 and F2 (Supplementary Material 1). Thus, to reduce the number of variables, a principal component analysis (PCA) was performed. The first axis of this PCA that well summarized the variability information contained in the two variables (PC1 > 95%; Supplementary Material 1) was interpreted as an estimation of snail size. The coordinate on the first PCA axis was extracted for all individuals (juvenile and adult stages separately) and analysed in two linear models (LM; one for the juvenile stage and one for the adult stage) using an analysis of variance (ANOVA) with predator exposure (factor variable with two levels), resource availability (factor variable with two levels) and their interaction. To give a biological estimate of the effects, the raw length and weight (and not the first PCA axis coordinate) were used to calculate the percentage of differences between means of environmental treatment. The figures of raw length and weight and the analyses are available in Supplementary Material 2. ***Crawling-out behaviour*** was a binary variable (0 or 1) and was then analysed with a Generalized Linear Model (GLM) using a binomial distribution (logit link function) and with the same predictors as for models of snail size. The significance of effects was tested using likelihood ratio tests. ***Shell crush resistance*** was analysed in a LM using an ANOVA with the same predictors as the other models. ***Reproductive outputs:*** the numbers of eggs and juveniles were analysed with a GLM using a negative binomial distribution to account for overdispersion and with the same predictors at the other models. The significance of effects was tested using likelihood ratio tests. Post-hoc pairwise comparisons (using a Tukey test) were performed on all models to identify differences between the combinations of environmental treatments. The versions of R and RStudio, as well as the R packages used and their respective versions, are detailed in the ReadMe file provided at the data and code repository.

## Results

### Size of F1 snails at juvenile and adult stages

F1 snail size at juvenile and adult stages were significantly influenced by the interaction between predator-cue exposure and resource availability (Table 1, Fig. 2). In the high resource environment, juvenile and adult snails exposed to predator cues were smaller than non-exposed snails (juvenile PH-CH: 12% shorter and 29% lighter, t[432] = -7.85, p < 0.001; adult PH-CH: 14% shorter and 36% lighter, t[432] = -14.47, p < 0.001). In the low resource environment, juvenile exposed to predator cues had a similar size than non-exposed snails (PL-CL: t[432] = -1.91, p = 0.221) but adult snails exposed to predator cues were smaller than non-exposed snails (PL-CL: 7% shorter and 19% lighter, t[432] = -3.58, p = 0.002). Overall, juvenile and adult snails reared in low resource were significantly smaller than those in high resource environment (Fig. 2; juvenile L-H: 40% shorter and 77% lighter; adult L-H: 40% shorter and 79% lighter).

**Figure 2.**
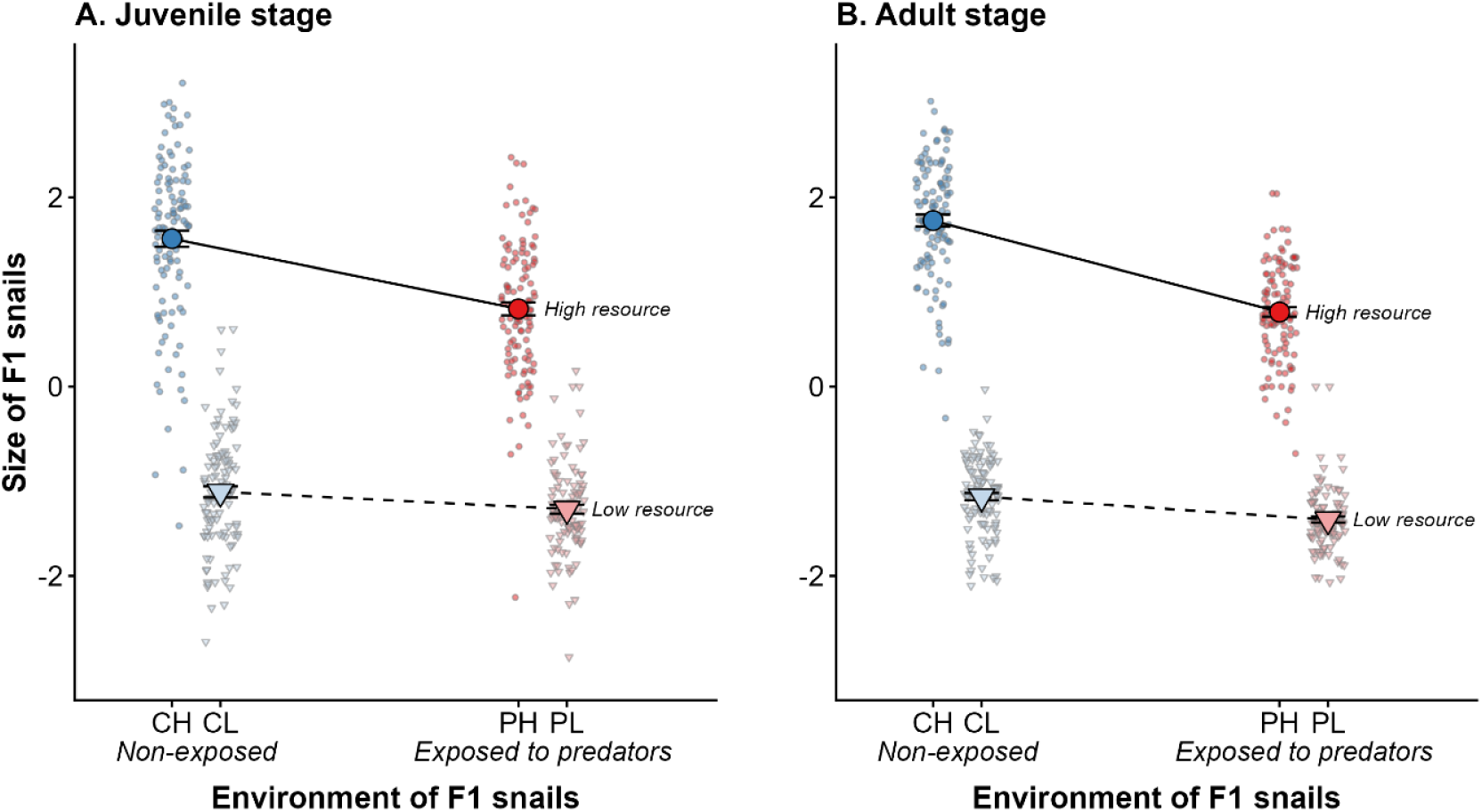
Effects of exposure to predator cues (Exposed P in red and non-exposed control C in blue) and resource availability (High H in bright colour/circle symbol and Low L in pale colour/triangle symbol) on the F1’s snail size at A-juvenile and B-adult stages. The solid and dashed lines represent the mean reaction norms in response to predator-cue exposure in high and low resource environment, respectively. Big and coloured dots are mean of raw data with their standard errors while small grey dots are individual values. Figures on total mass and shell length are given in Supplementary Material 2.

**Table 1:** Analyses of variance testing the effects of exposure to predator cues and resource availability on traits of F1 snails. Defined intercepts are control non-exposed to predator-cue (C) and high resource (H) environments. F-values or X² statistics are shown according to the models. Bold P-values are significant (< 0.05).

| Trait | Effect | Estimate [SE] | Effect size $\eta^2$ [CI] | NumDf , DenDf | $X^2$ or F-value | P-value |
| --- | --- | --- | --- | --- | --- | --- |
| Juvenile size | Predation | -0.74 [0.09] | 0.09 [0.05, 1] | 1, 432 | 47.81 | **< 0.001** |
|  | Resource | -2.67 [0.09] | 0.75 [0.72, 1] | 1, 432 | 1281.20 | **< 0.001** |
|  | P:R | 0.56 [0.13] | 0.04 [0.01, 1] | 1, 432 | 17.49 | **< 0.001** |
| Adult size | Predation | -0.97 [0.07] | 0.25 [0.20, 1] | 1, 432 | 163.49 | **< 0.001** |
|  | Resource | -2.92 [0.07] | 0.87 [0.86, 1] | 1, 432 | 2921.31 | **< 0.001** |
|  | P:R | 0.73 [0.09] | 0.12 [0.08, 1] | 1, 432 | 58.80 | **< 0.001** |
| Crawling-out behaviour | Predation | 2.40 [0.33] |  | Df = 1, N = 424 | 191.89 | **< 0.001** |
|  | Resource | -0.85 [0.38] |  | Df = 1, N = 424 | 0.05 | 0.828 |
|  | P:R | 1.53 [0.53] |  | Df = 1, N = 424 | 8.71 | **0.003** |
| Shell crush resistance | Predation | 0.07 [0.02] | 0.02 [0.00, 1] | 1, 407 | 8.54 | **< 0.01** |
|  | Resource | -0.02 [0.02] | 0.01 [0.00, 1] | 1, 407 | 5.95 | **0.01** |
|  | P:R | -0.04 [0.03] | < 0.01 [0.00, 1] | 1, 407 | 1.18 | 0.278 |
| Egg number | Predation | -0.37 [0.17] |  | Df = 1, N = 419 | 2.82 | 0.0930 |
|  | Resource | -2.01 [0.17] |  | Df = 1, N = 419 | 209.39 | **< 0.001** |
|  | P:R | 0.36 [0.24] |  | Df = 1, N = 419 | 2.21 | 0.137 |
| Juvenile number | Predation | -0.31 [0.16] |  | Df = 1, N = 419 | 1.38 | 0.24 |
|  | Resource | -1.96 [0.16] |  | Df = 1, N = 419 | 208.11 | **< 0.001** |
|  | P:R | 0.35 [0.23] |  | Df = 1, N = 419 | 2.24 | 0.135 |

### Anti-predator behaviour and shell resistance of F1 adult snails

The number of adult F1 snails exhibiting the crawling-out behaviour was influenced by the interaction between predator-cue exposure and resource availability (Table 1, Fig. 3A). Predator cues led to an increase in crawling-out behaviour in both resource environments (PH-CH: odds.ratio = 11.07, z = 7.38, p < 0.001; PL-CL: odds.ratio = 51.39, z = 9.38, p < 0.001). This increase is slightly higher for low resource snails. However, the low resource availability did not significantly influence the behaviour both in non-exposed individuals (CL-CH: odds.ratio = 0.43, z = -2.21, p = 0.120) and in predator-cue exposed ones (PL-PH: odds.ratio = 1.99, z = 1.87, p = 0.242) relative to the high resource environment.

**Figure 3:**
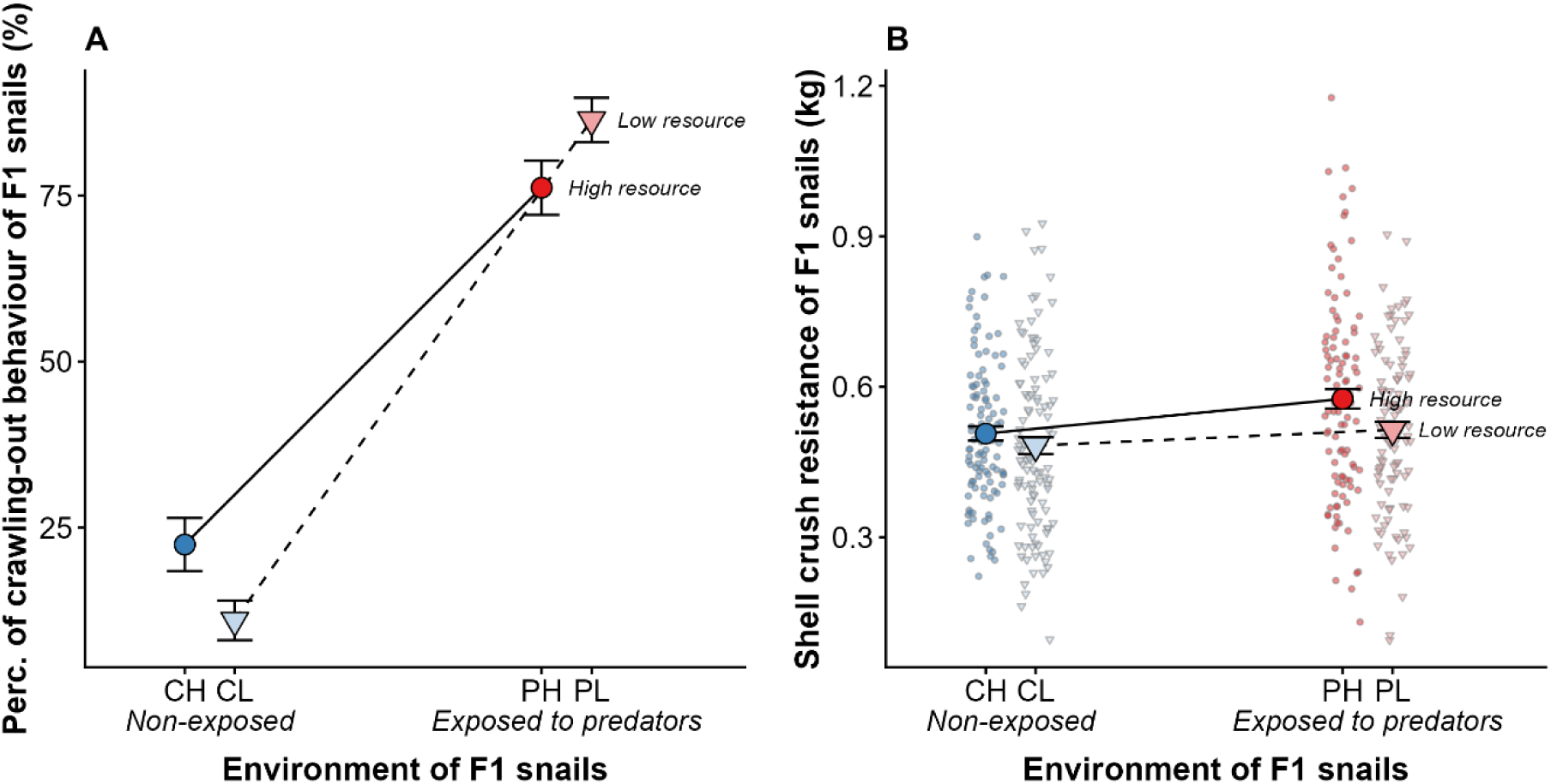
Effects of exposure to predator cues (Exposed P in red and non-exposed control C in blue) and resource availability (High H in bright/circle symbol colour and Low L in pale colour/triangle symbol) on F1’s A-crawling-out behaviour and B-shell crush resistance. The solid and dashed lines represent the mean reaction norms in response to predator-cue exposure in high and low resource environment, respectively. Big and coloured dots are mean of raw data with their standard errors while small grey dots are individual values.

Shell crush resistance was independently influenced by predator-cues exposure and resource availability (non-significant interaction, Table 1, Fig. 3B). Shell crush resistance was significantly 10% higher in snails exposed to predator-cues relative to non-exposed snails regardless the resource environment and was 7.9% lower in low resource compared to high resource regardless the predator exposure (non–significant interaction, Table 1, Fig. 3B).

### Reproductive outputs of F1 adult snails

Egg and juvenile numbers produced by the F1 adult snails were only influenced by resource availability (Table 1, Fig. 4). Snails raised in low resource laid significantly fewer eggs and produced fewer juveniles than those raised in high resource (-84% and -83%, respectively).

**Figure 4:**
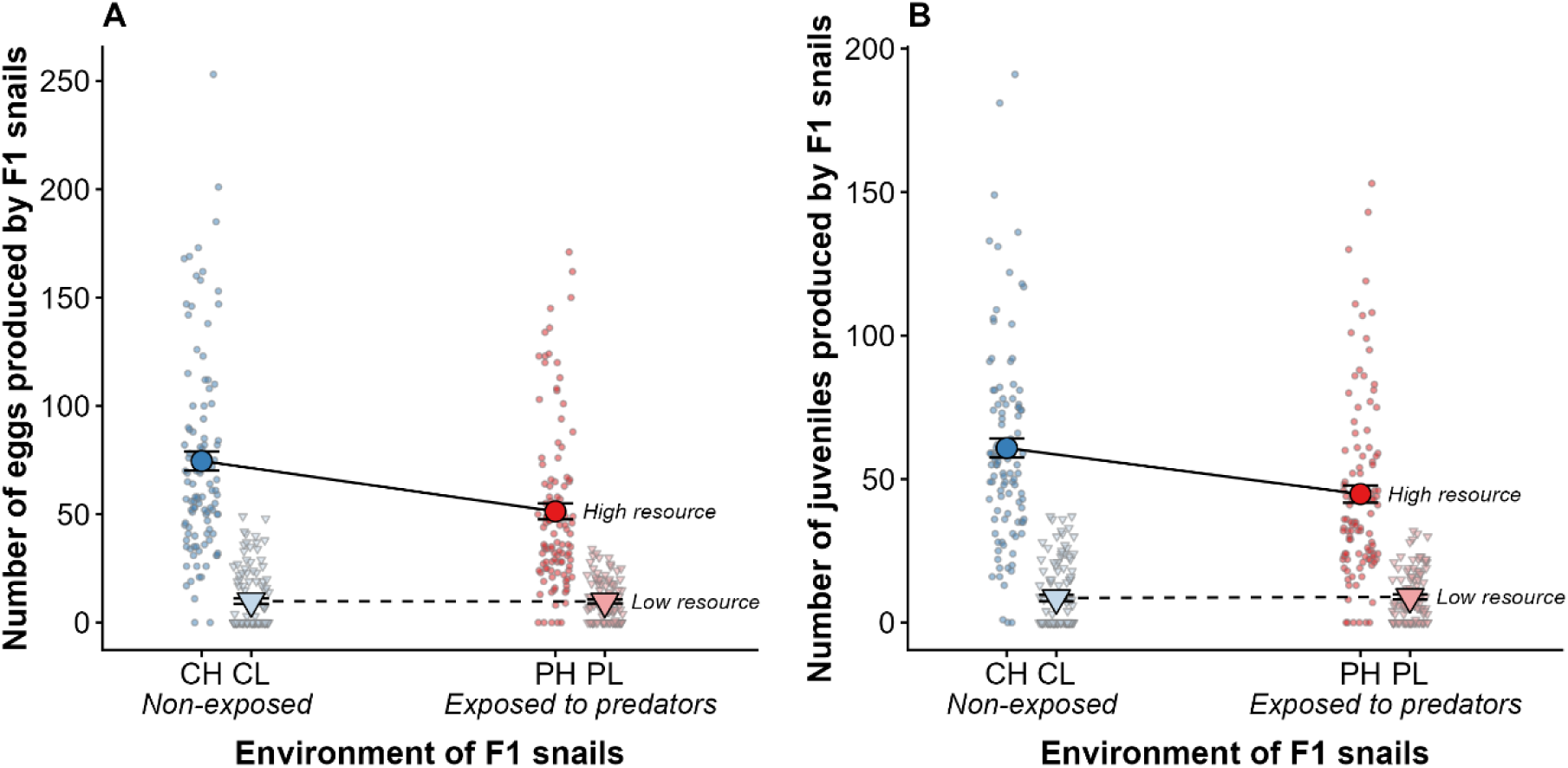
Effects of exposure to predator cues (Exposed P in red and non-exposed control C in blue) and resource availability (High H in bright colour/circle symbol and Low L in pale colour/triangle symbol) on the number of A-eggs and B-juveniles produced by the F1 snails. The solid and dashed lines represent the mean reaction norms in response to predator-cue exposure in high and low resource environment, respectively. Big and coloured dots are mean of raw data with their standard errors while small grey dots are individual values.

### Size of F2 snails at juvenile and adult stages

F2 snail size at the juvenile and adult stages were significantly influenced by the interaction between parental predator-cue exposure and parental resource availability (interaction is marginally significant for adults, see Table 2, Fig. 5). The parental exposure to predator-cues significantly increased the juvenile and adult size of F2 snails from both resource parental environment (juvenile PH-CH: 9% longer and 3% heavier, t[444] = 2.73, p = 0.033; juvenile PL-CL: 35% longer and 92% heavier, t[444] = 8.66, p < 0.001; adult PH-CH: 3% longer and 12% heavier, t[444] = 3.17, p = 0.010; adult PL-CL: 7% longer and 26% heavier, t[444] = 5.60, p < 0.001); this increase was slightly higher for F2 snails from low resource parental environment (Table 2, Fig. 5). Overall, F2 juvenile and adult snails from the low resource parental environment were significantly smaller than F2 ones from the high resource parental environment (Fig. 5; juvenile CL-CH: 41% shorter and 80% lighter, t[444] = -24.85, p < 0.001; juvenile PL-PH: 32% shorter and 62% lighter, t[444] = -18.91, p < 0.001; adult CL-CH: 8% shorter and 27% heavier, t[444] = -7.51, p < 0.001; adult PL-PH: 4% shorter and 18% lighter, t[444] = -5.11, p < 0.001).

**Figure 5:**
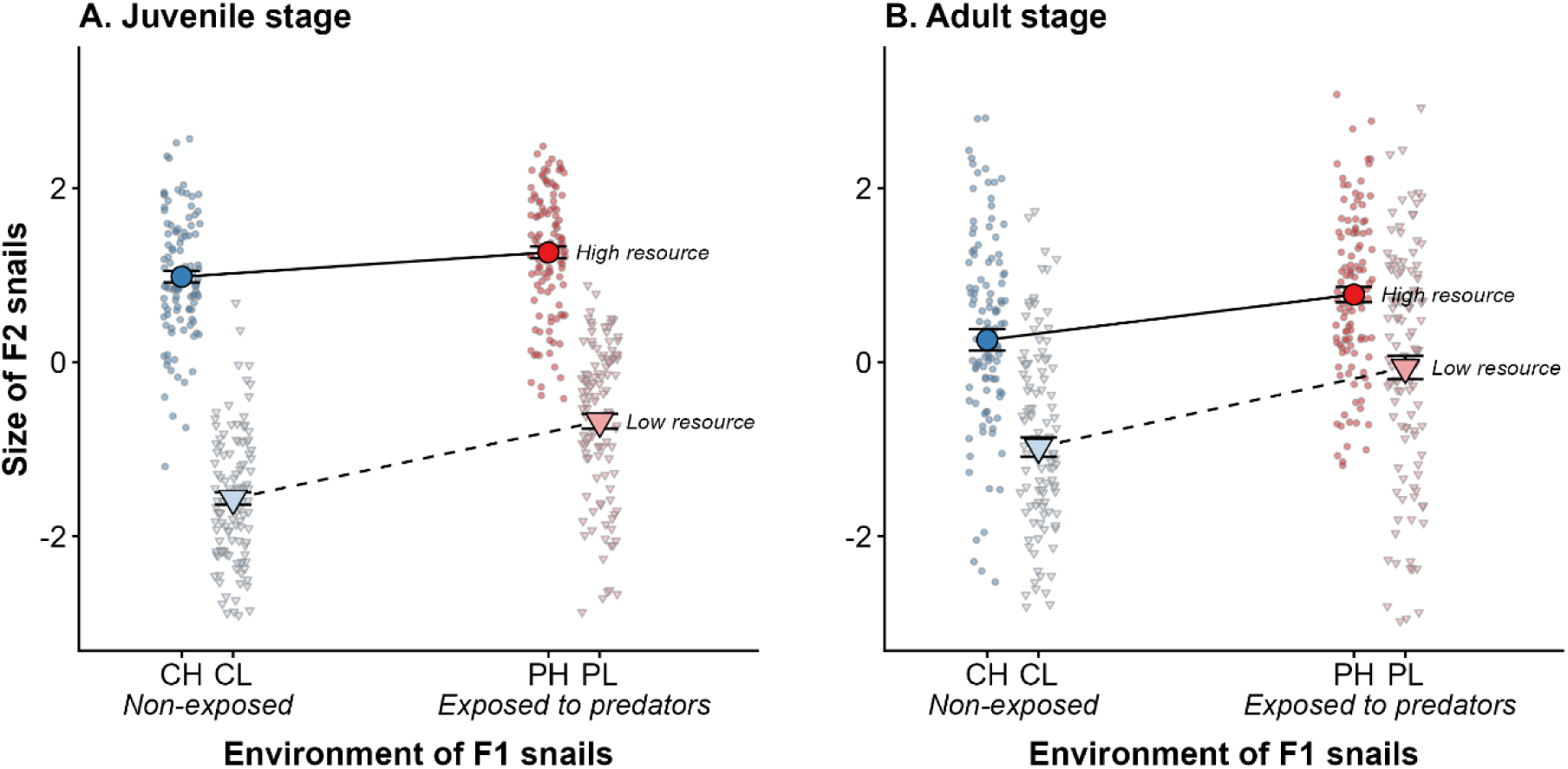
Effects of parental exposure to predator cues (Exposed P in red and non-exposed control C in blue) and resource availability (High H in bright colour/circle symbol and Low L in pale colour/triangle symbol) on offspring size at the A-juvenile and B-adult stages. The solid and dashed lines represent the mean transgenerational reaction norms in response to parental predator-cue exposure in high and low parental resource environment, respectively. Big and coloured dots are mean of raw data with their standard errors while small grey dots are individual values. Figures on total mass and shell length are given in supplementary material 2.

**Table 2:** Analyses of variance testing the effects of exposure to predator cues and resource availability on traits of F2 snails. Defined intercepts are control non-exposed to predator-cue (C) and high resource (H) environments. F-values or X² statistics are shown according to the models. Bold P-values are significant (< 0.05).

| Trait | Effect | Estimate [SE] | Effect size $\eta^2$[CI] | NumDf , DenDf | $\chi^2$ or F-value | P-value |
| --- | --- | --- | --- | --- | --- | --- |
| Juvenile size | Predation | 0.28 [0.10] | 0.13 [0.08, 1] | 1, 444 | 64.91 | **< 0.001** |
|  | Resource | -2.55 [0.10] | 0.68 [0.65, 1] | 1, 444 | 957.46 | **< 0.001** |
|  | P:R | 0.61 [0.14] | 0.04 [0.01, 1] | 1, 444 | 17.61 | **< 0.001** |
| Adult size | Predation | 0.52 [0.16] | 0.08 [0.04, 1] | 1, 444 | 38.13 | **< 0.001** |
|  | Resource | -1.23 [0.16] | 0.15 [0.10, 1] | 1, 444 | 79.67 | **< 0.001** |
|  | P:R | 0.39 [0.23] | < 0.01 [0.00, 1] | 1, 444 | 2.88 | 0.090 |
| Crawling-out behaviour | Predation | 1.25 [0.29] |  | Df = 1, N = 438 | 18.85 | **< 0.001** |
|  | Resource | 0.49 [0.29] |  | Df = 1, N = 438 | 0.16 | 0.686 |
|  | P:R | -0.77 [0.40] |  | Df = 1, N = 438 | 3.77 | 0.052 |
| Shell crush resistance | Predation | 0.01 [0.02] | < 0.01 [0.00, 1] | 1, 426 | 4.42 | **0.036** |
|  | Resource | -0.10 [0.02] | 0.09 [0.05, 1] | 1, 426 | 41.08 | **< 0.001** |
|  | P:R | 0.03 [0.03] | < 0.01 [0.00, 1] | 1, 426 | 1.49 | 0.22 |
| Egg number | Predation | -0.08 [0.07] |  | Df = 1, N = 435 | 2.01 | 0.148 |
|  | Resource | -0.27 [0.07] |  | Df = 1, N = 435 | 13.52 | **< 0.001** |
|  | P:R | 0.16 [0.10] |  | Df = 1, N = 435 | 2.61 | 0.106 |
| Juvenile number | Predation | -0.01 [0.08] |  | Df = 1, N = 435 | 4.27 | **0.038** |
|  | Resource | 0.17 [0.08] |  | Df = 1, N = 435 | 24.12 | **< 0.001** |
|  | P:R | 0.19 [0.11] |  | Df = 1, N = 435 | 3.21 | 0.072 |

### Anti-predator behaviour and shell resistance of F2 adult snails

The number of F2 adult snails exhibiting the crawling-out behaviour was marginally influenced by the interaction between parental resource and parental exposure to predator-cues (Table 2, Fig. 6A). Parental exposure to predator cues increased the crawling-out behaviour of F2 adult snails in the high resource parental environment (PH-CH: odds.ratio = 3.51, z = 4.30, p = 0.001) but did not significantly affect their behaviour in the low resource parental environment (PL-CL: odds.ratio = 1.62, z = 1.76, p = 0.293). The parental resource environment did not significantly affect the offspring behaviour from non-exposed parents (CL-CH: odds.ratio = 1.64, z = 1.68, p = 0.333) and exposed parents (PL-PH: odds.ratio = 0.75, z = -1.03, p = 0.730).

**Figure 6:**
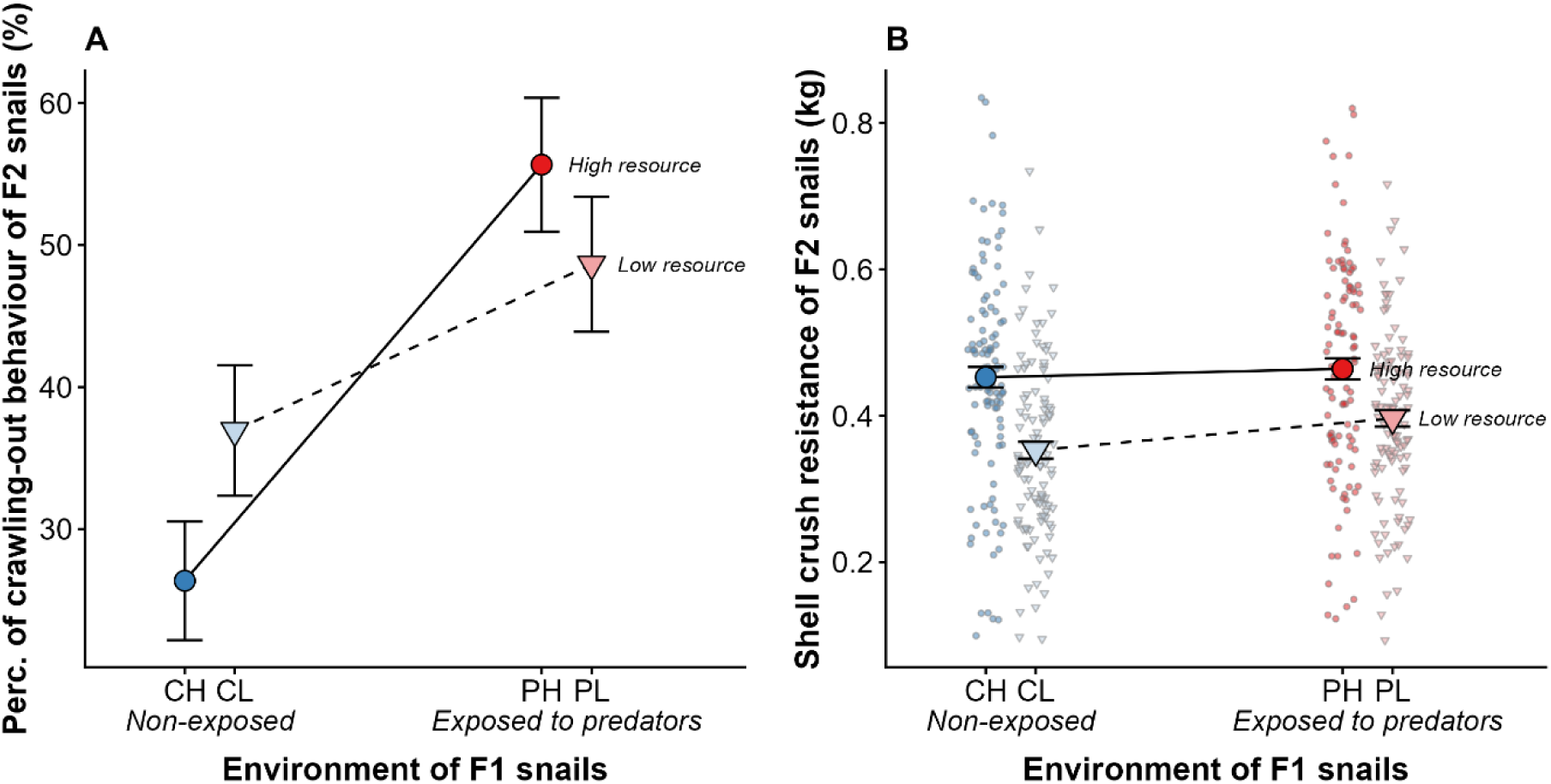
Effects of parental exposure to predator cues (Exposed P in red and non-exposed control C in blue) and resource availability (High H in bright colour/circle symbol and Low L in pale colour/triangle symbol) on F2 offspring A- crawling-out behaviour and B- shell crush resistance. The solid and dashed lines represent the mean transgenerational reaction norms in response to parental predator-cue exposure in high and low parental resource environment, respectively. Big and coloured dots are mean of raw data with their standard errors while small grey dots are individual values.

Shell crush resistance of F2 snails was independently influenced by the parental resource and the parental exposure to predator-cues (non-significant interaction, Table 2, Fig. 6B). The shell resistance increased (+7%) for F2 snails from parents exposed to predator-cues relative to those from non-exposed parents. Shell was 18% less resistant for F2 snails from the low resource parents.

### Reproductive outputs of F2 adult snails

Egg number laid and juvenile number produced by F2 snails were influenced by the parental resource (interaction with parental predator-cue exposure marginally significant for juvenile number; Table 2, Fig. 7). F2 snails from the low resource parental environment produced fewer eggs (−17%) but more juveniles (+31%) than those from the high resource parental environment (Fig. 7). In addition, F2 snails from exposed parents produce more juveniles (+12%) than those from non-exposed parents (Table 2; Fig. 7B). In the high resource parental environment, there were no difference between F2 snails from exposed and non-exposed parents (egg number PH-CH: ratio = 0.99, z = -0.11, p = 0.999; juvenile number PH-CH: ratio = 1.01, z = -0.19, p = 0.997). In the low resource parental environment, F2 snails from exposed parents laid a similar number of eggs but produced more juvenile than those from non-exposed parents (egg number PL-CL: ratio = 1.17, z = 2.17, p = 0.132; juvenile number PL-CL: ratio = 1.23, z = 2.73, p = 0.032).

**Figure 7:**
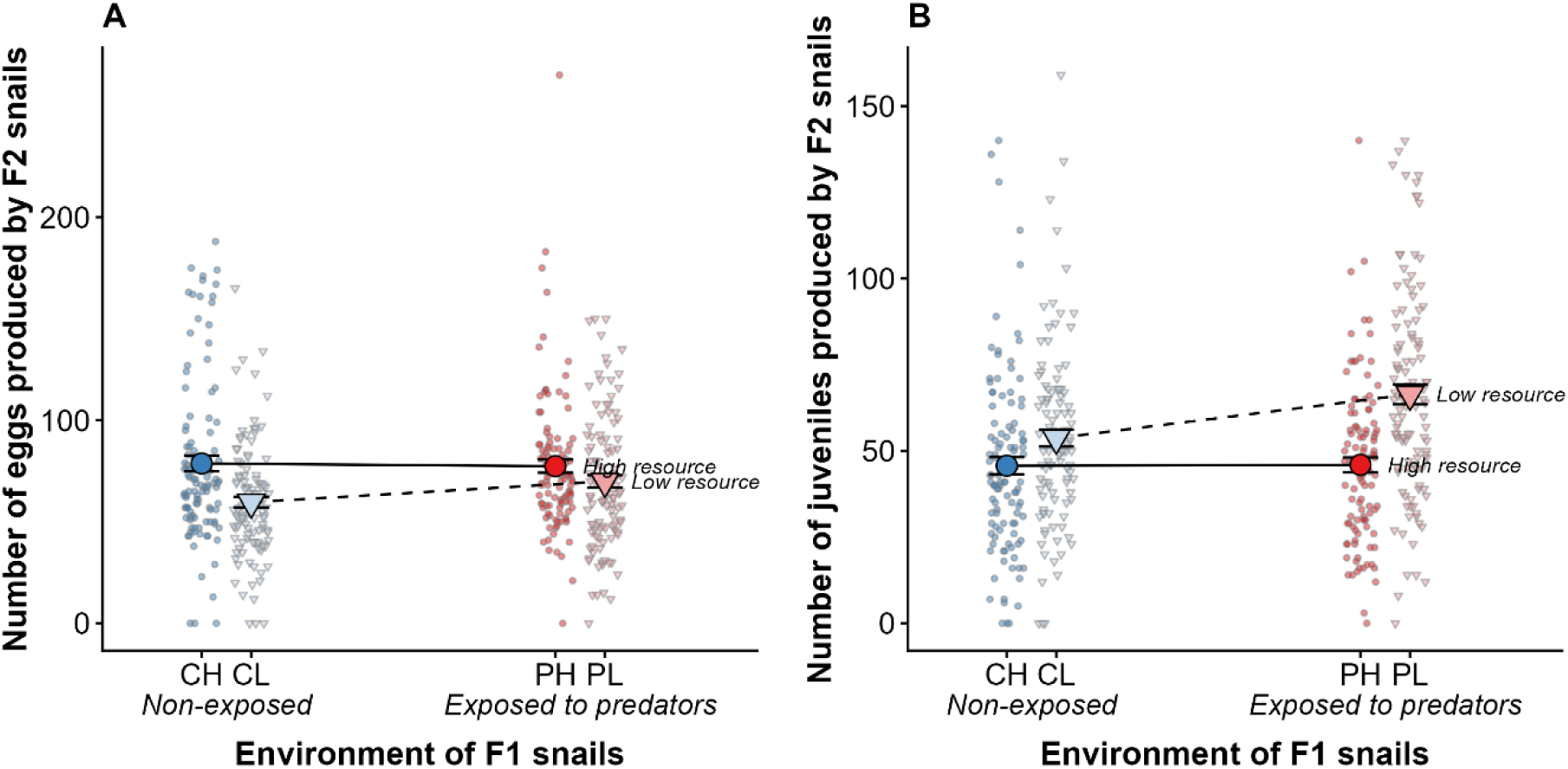
Effects of parental exposure to predator cues (Exposed P in red and non-exposed control C in blue) and resource availability (High H in bright colour/circle symbol and Low L in pale colour/triangle symbol) on the number of A- eggs and B- juveniles produced by F2 offspring. The solid and dashed lines represent the mean transgenerational reaction norms in response to parental predator-cue exposure in high and low parental resource environment, respectively. Big and coloured dots are mean of raw data with their standard errors while small grey dots are individual values.

## Discussion

Phenotypic plasticity, both WGP and TGP, can occur in response to environmental information (cues) that predict the future (selective) environment or as a consequence of environmental alteration on the individual’s somatic state [15]. In the context of predator-prey interactions, prey face a trade-off between developing traits beneficial for acquiring resources or for defending against predators, closely intertwining information- and somatic state-based effects. In this study, our aim was to show how these types of plasticity can interact to shape the phenotypic responses by manipulating the resource acquisition-safety trade-off. Prerequisites to investigate such an interaction were to provide evidence of information- and state-based WGP and TGP. As expected, our results revealed that F1 exposure to predator cues triggered changes in anti-predator traits and growth of F1 and F2 individuals, illustrating information-based WGP and TGP (see consistent previous studies on freshwater snails e.g. [34,35,45–47] or in other taxa e.g. [48,49]). Our study also confirmed that developmental exposure of F1 individuals to limited resource availability constrained growth in both F1 and F2 generations. F1 individuals reared in a limited-resource environment globally reached a smaller size, indicating a poorer somatic state of individuals and illustrating state-based WGP (see consistent examples in freshwater snails e.g. [10,46,50,51] or in other taxa e.g. [52,53]). The poorer F1 parental state in low-resource environment had a carry-over effect on F2 offspring state (i.e. F2 individuals from low-resource parents were smaller and lighter than those from high-resource parents), illustrating state-based TGP (see reviews [23,54] and an example on *Daphnia* [53]). In the following, we discuss the results revealing that information- and state-based plasticity combined to shape additive and non-additive patterns of WGP and TGP reaction norms through the resource acquisition-safety trade-off.

### The combination of information- and state-based plasticity shapes the WGP reaction norms

Developmental exposure to predator cues is well known to induce information-based WGP (28), including shifts in life-history traits (e.g. growth, fecundity) as well as specific behavioural and morphological anti-predator traits, across many organisms ranging for example from birds [55], lizards [56] and amphibians [57] to molluscs [47], insects [58] and crustacean [59]. Changes in life-history traits may arise either as direct responses to predation cues (e.g. they can be defensive) or as by- products of the induction of behavioural and morphological defences through energetic trade-offs, thereby intertwining state- and information-based plasticity [27]. For instance, Tollrian (1995) showed, in *Daphnia pulex*, that the life history shifts in response to predator cues were independent of morphological changes (“neckteeth”)[60]. Conversely, Teplitsky et al. (2003) demonstrated that the induction of morphological defences in tadpoles of *Rana dalmatina* was associated with reduced growth and larval development rates, reflecting potential energetic costs of producing anti-predator traits [61].

In the same way, our results revealed the induction of anti-predator traits – namely increased crawling-out behaviour and shell-crush resistance – in response to predator cues associated with a reduced growth, consistently with our previous studies (e.g. [36,42,45]). The reduced individual size is a non-consumptive effect commonly reported under predation risk in freshwater snails ([62–64]; but see [3,34,65]), and reflects energetic trade-offs behind information-based WGP of defences [27]. By experimentally manipulating resource availability and predator cue exposure, our study also provided evidence that the WGP reaction norms were shaped by a combination of state- and information-based plasticity:

First, the reduction in individual size in response to predator-cues observed in high-resource environment, was in low-resource environment absent (juveniles) or less pronounced (adults), revealing a masking non-additive pattern between the state- and information-based plasticity. The smaller baseline size of snails exposed to limited resources may impose physical constraints that limit further reduction due to a reallocation of energy away from growth. For instance, other studies on freshwater snails and amphibians found such a masking non-additive pattern by revealing that the negative effects of predation risk on growth were strongest when prey are at low density (i.e. low competition for resources) but weaken or disappear completely when prey are at high density (i.e. high competition for resources; [10,66–69]).

Second, despite resource limitations, individuals from the low-resource environment still expressed the anti-predator traits (crawling-out behaviour, shell-crush resistance) at levels comparable to those from the high-resource environment. Interestingly, a revealing non-additive pattern was observed for crawling-out behaviour: the slope of the reaction norm increased for individuals from low-resource environment, a pattern already observed in another *Physa* species [70]. This may suggest a compensation for heightened vulnerability to crayfish predation due to a smaller size in low resource environment by an increase of crawling-out behaviour. Such trait compensation in prey facing predators has been observed in several studies on freshwater snails (e.g. [71–73]) and also for example on birds [74] and fishes [75,76]. In contrast, we found no interaction between information- and state-based plasticity in shell-crush resistance (i.e. additive pattern). The shell crush resistance increased with predator-cues exposure with the same magnitude in individuals from low- and high resource environments.

Third, the exposure to predator cues did not influence the reproductive outputs but the low-resource environment strongly reduced it. This suggests that individuals exposed to predator cues still find the resources they need to invest in reproduction, even in low-resource environment. Brönmark et al. (2012) [77] found consistently that defence traits were expressed independent of environmental quality (individual density) in the freshwater snail *Radix balthica* and resulted in reduced growth rate but also decreased fecundity, particularly with limited resources (see also [46]).

Overall, our results demonstrate that exposure to predator cues influence the somatic state, but above all, even under resource scarcity, the induction of anti-predator traits remains a priority, drawing energy away from growth (but not from reproduction), which suggests strong selection from predators. Such investment in defences might improve individual fitness by improving survival under predation risk and thus by extending lifespan, thereby allowing reduced reproductive output, in particular in low-resource environment, to be compensated across multiple reproductive events [78]. This echoes the general prediction that selection maximizes parental fitness rather than offspring fitness, especially in species that lack parental care and that reproduce many times [26]. Additional research evaluating lifetime reproductive success of *P. acuta* under predation risk and resource scarcity must be conducted before definite conclusions about the adaptive value of phenotypic responses observed.

### The combination of information- and state-based plasticity shapes the TGP reaction norms

Our results provided evidence that, overall, the parental environments influenced the offspring traits, indicating TGP as previously observed in this system (e.g. [36,39,45]). Parental exposure to predator cues increased the size, crawling-out behaviour and shell-crush resistance of F2 individuals. This suggests adaptive transgenerational information-based responses, whereby parental information ‘anticipates’ the risk of predation that offspring will experience and to induce the defences of offspring accordingly [17,26]. Indeed, increased size might enable individuals to reach a “refuge size”, reducing vulnerability to crayfish predation (as crayfish preferentially consume smaller snails; [3,34,71]), while enhanced crawling-out behaviour and shell crush resistance—resulting from a complex combination of shifts in shell size, shape, and thickness—improves the likelihood of surviving predator attacks [35,39,79,80]. Additional work is nevertheless necessary to test the adaptive value of transgenerational responses of *P. acuta* to crayfish.

Above all, our results demonstrated that the combination of parental resource availability and exposure to predator cues shaped the TGP reaction norms via an additive pattern for shell crush resistance and non-additive masking or revealing patterns for size, crawling-out behaviour and marginally for reproductive outputs. The overall weaker shell crush resistance of F2 from low-resource parents may be a physical consequence of their smaller size compared to F2 from high-resource. By constraining the absolute level of size, state-based TGP influenced in turn the absolute level of shell crush resistance in absence of actual predator cues, which might influence the predator-induced WGP responses (not tested in this study but see [36]). The increase in size in response to parental predator cue exposure was slightly higher for F2 snails from low-resource parents (revealing pattern), suggesting that these individuals may take advantage of their favourable high resource current environment to partly compensate for the state-based TGP constraining their size. In contrast, crawling-out behaviour of F2 individuals exhibited a masking pattern: parental exposure to predator cues induced crawling-out only in F2 from high-resource parents, while no such response was observed in F2 from low-resource parents. The lack of significant behavioural response to parental predator exposure in F2 from low-resource parents might illustrate the trade-off between resource acquisition and anti-predator traits. Indeed, a poor state-driven effect may cause an elevated baseline crawling-out behaviour in control individuals and a lower crawling-out behaviour in parental exposed individuals. The elevated baseline crawling-out behaviour might reflect a behavioural compensation for morphological vulnerability (i.e. small size) even in the absence of parental and actual predator exposure [71]. The poor state of F2 from low-resource parents might also constraint the time spent to crawl-out at the expense of safety to prioritize foraging in the water. For example, Relyea (2004) showed that increased competition (i.e. low resource condition) caused higher activity of *Rana sylvatica* tadpoles, even under predation risk[66]. These results illustrate well that the intertwining between information- and state-based effects can drive TGP reaction norms through the resource acquisition-safety trade-off: current environmental conditions, such as the risk of predation, can be perceived by parents in order to predict the future environment of their offspring and adjust their phenotype accordingly, but these anticipatory responses may be constrained because such environmental conditions also influence the body state of the parents and, consequently, that of their offspring [16,18,23,53].

Finally, the results may suggest a revealing pattern for juvenile number: parental exposure to predator cues increased the numbers of juveniles only in F2 snails from low-resource parents. We further showed that this predator-induced TGP on juvenile number was mediated by offspring size, since the effect of parental exposure to predator cues disappeared once mass was accounted for (Supplementary Material 3). The positive relationship between size and reproductive outputs is well documented in *P. acuta* (e.g. [78,81]) and in ectotherms in general ([82]). Consequently, the increased juvenile number produced by F2 individuals from low-resource and predator-exposed parents may not reflect an energy reallocation from growth to reproduction, but rather a by-product of changes in size and defences driven by the interaction between parental state and parental predator information. Interestingly, F2 snails from low-resource and predator-exposed parents exhibited the highest juvenile number overall. While they produced a similar number of eggs as snails from high-resource and predator-exposed parents, they produced a greater number of surviving juveniles. This suggests a higher hatching success and early juvenile survival (for 3 days post-hatching) in the offspring (F3) of F2 from low-resource and predator-exposed parents. Thus, F2 individuals may improve their overall reproductive success (by increasing F3 offspring fitness) in absence of current predator cues under combined parental state- and information-based effects by modulating the trade-off between offspring number and offspring quality. This result is rare evidence that fitness benefits of TGP are condition-dependent (here a combination of ancestral exposure to predator cues and low resource availability) and can be revealed only three-generations later (see evidence for grand-parental effects in *P. acuta* [42] or in other taxa e.g. [83–85]), extending the widely documented condition-dependent parental effects on reproductive strategy in many organisms (see reviews [23,25,26]). Further investigation of maternal provisioning in eggs (e.g. egg size, energetic reserves in eggs) would be valuable to unravel the proximal mechanisms behind the transgenerational modulation of the trade-off between offspring number and offspring quality.

## Conclusion

Our study has shown that both information- and state-based plasticity participated in shaping the phenotypic responses through the resource acquisition-safety trade-off according to additive and non-additive patterns within and throughout generations. Within a generation, the state-based WGP did not constrain the costly expression of inducible defences (information-based WGP), although it decreased the individual’s somatic state, suggesting a strong selective pressure from predation. The expression of anti-predator traits was associated with costs influencing the individual’s somatic state; costs that could be paid via different trade-offs according to the resource availability. Across generations, the poor state of parents significantly decreased the state of offspring, which in turn influenced the levels of their defences. These results illustrate well the intertwining between information- and state-based effects that can drive TGP patterns. While the offspring somatic state determined from that of their parents constrained the offspring responses to parental information, the parental information prepared offspring for both predation risk and resource limitation, buffering the parental state-based effect. Note however that the higher vulnerability to predation of offspring from low resource parents and a possible buffering effect by the higher reproductive success must been addressed in a context of predator exposure to draw definite conclusions. Indeed, in our experimental conditions the state-based TGP may have had less impact on the predator-induced TGP because the offspring (F2 individuals) were facing extremely favourable conditions throughout their life (“beneficially saturated” conditions) while harsher conditions would have amplified or saturated the effect of parental condition (“detrimentally saturated” conditions; [18]). Finally, our study, by integrating both information and state-based plasticity across generations, provides a more complete understanding of prey responses to predators according to resource availability and, in general, important information regarding the patterns of phenotypic integration across environments and generations.

## Acknowledgements

We are grateful to Pierre Grancha, Anouk Piednoir, Chloé Marmonier and Théodora Icarre who helped in experimental works. We are also grateful to Ludovic Guillard who designed and built the electronic device that measures the shell crush resistance.

## Authors Contributions

L.D., S.P. and E.L. conceived the project. L.D., S.P. and E.L. performed the experiments. L.D. and J.T.A. analysed the data. L.D. wrote the first version of the manuscript. All authors revised the manuscript.

## Ethics

This work did not require ethical approval from a human subject or animal welfare committee.

