## Supplementary material for "Resource availability and predator cues shape within- and transgenerational reaction norms"

### Supplementary materials

**Supplementary material 1:** Snail size of F1 and F2 – PCA on shell length and total mass

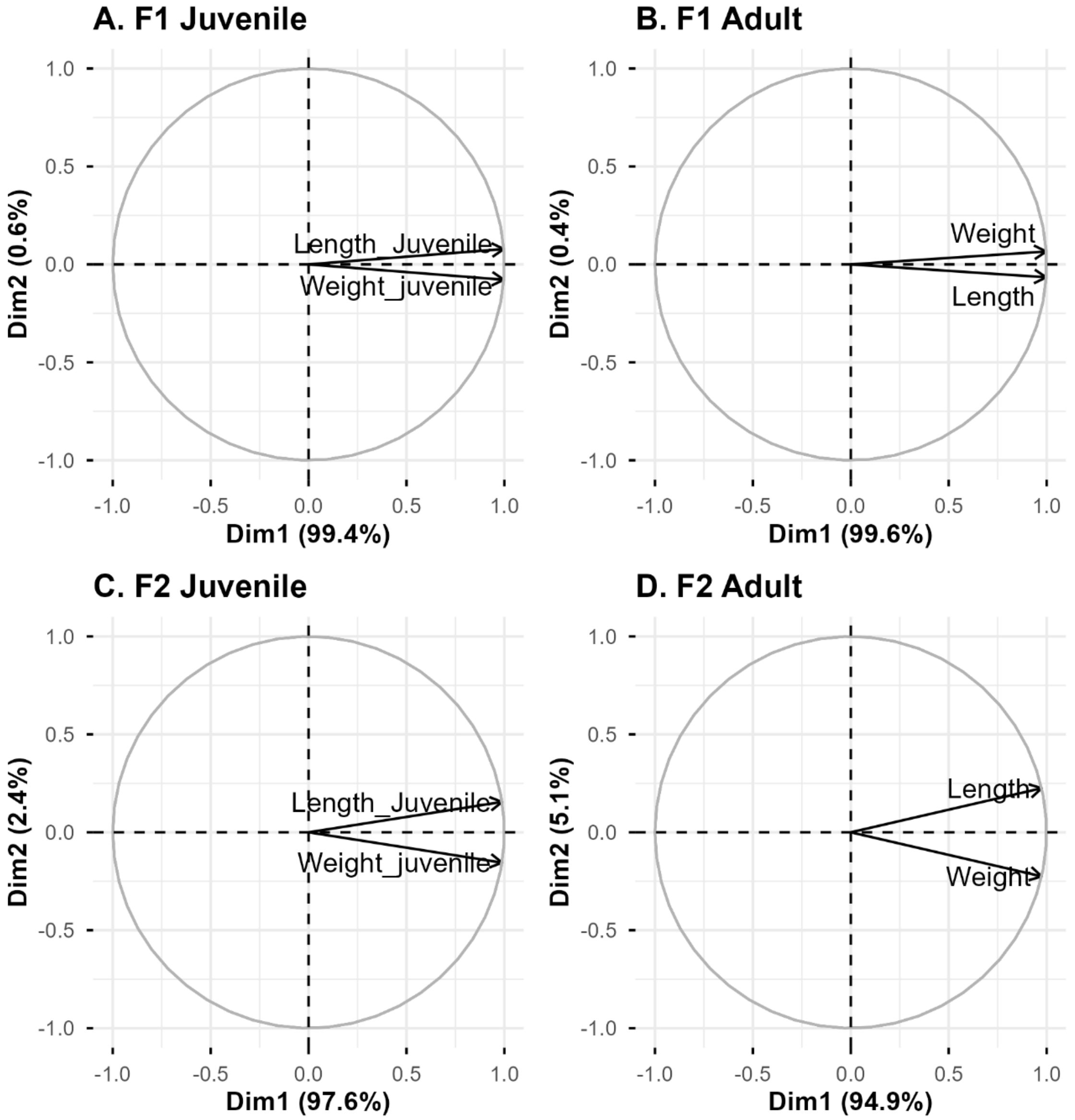


Correlation circles of the Principal Component Analyses on A- F1 juvenile length and weight, B- F1 adult length and weight, C- F2 juvenile length and weight, and D- F2 adult length and weight.

**Supplementary material 2**

**A. F1 weight and length at juvenile and adult stages**

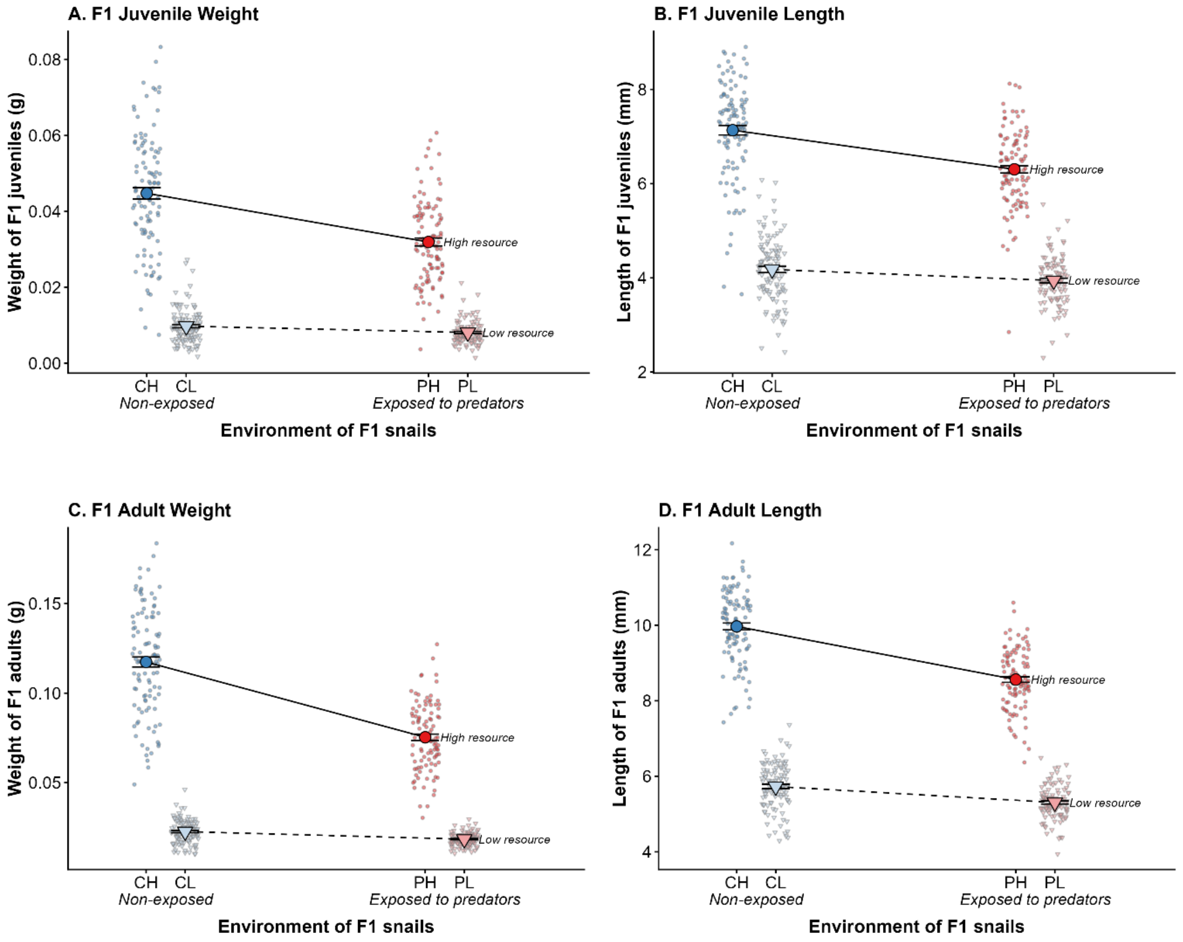


F1 snail weight and length at juvenile and adult stages were significantly influenced by the interaction between predator-cue exposure and resource availability (see Table below). In the high resource environment, juvenile and adult snails exposed to predator cues were shorter and lighter than non-exposed snails (juvenile PH-CH: 12% shorter, t[424] = -7.68, $p < 0.001$ and 29% lighter, t[422] = -9.44, $p < 0.001$; adult PH-CH: 14% shorter, t[420] = -14.14, $p < 0.001$ and 36% lighter, t[419] = -17.23, $p < 0.001$). In the low resource environment, juvenile exposed to predator cues had a similar length and weight than non-exposed snails (PL-CL: length t[424] = -2.17, p = 0.133; weight t[422] = -1.22, p = 0.615). Adult snails exposed to predator cues were shorter than non-exposed snails (PL-CL: 7% shorter, t[420] = -4.25, $p < 0.001$), but had a similar weight (t[419] = -1.73, p = 0.311). Overall, juvenile and adult snails reared in low resource were significantly shorter and lighter than snails in high resource environment (juvenile L-H: 40% shorter and 77% lighter; adult L-H: 40% shorter and 79% lighter).

| Trait | Effect | Estimate [SE] | Effect size $\eta^2$[CI] | NumDf , DenDf | F_value | P_value |
|---|---|---|---|---|---|---|
| **F1 Juvenile weight** | Predation | -0.01 [0.001] | 0.11 [0.06, 1] | 1, 422 | 56.88 | < **0.001** |
| | Resource | -0.35 [0.001] | 0.69 [0.66, 1] | 1, 422 | 947.48 | < **0.001** |
| | P:R | 0.01 [0.002] | 0.07 [0.04, 1] | 1, 422 | 33.79 | < **0.001** |
| **F1 Juvenile length** | Predation | -0.83 [0.11] | 0.09 [0.05, 1] | 1, 424 | 48.54 | < **0.001** |
| | Resource | -2.96 [0.10] | 0.74 [0.71, 1] | 1, 424 | 1214.51 | < **0.001** |
| | P:R | 0.59 [0.15] | 0.03 [0.01, 1] | 1, 424 | 15.17 | < **0.001** |
| **F1 Adult weight** | Predation | -0.04 [0.002] | 0.28 [0.23, 1] | 1, 419 | 180.04 | < **0.001** |
| | Resource | -0.09 [0.002] | 0.82 [0.80, 1] | 1, 419 | 1945.09 | < **0.001** |
| | P:R | 0.04 [0.003] | 0.22 [0.17, 1] | 1, 419 | 119.71 | < **0.001** |
| **F1 Adult length** | Predation | -1.41 [0.10] | 0.27 [0.21, 1] | 1, 420 | 169.06 | < **0.001** |
| | Resource | -4.24 [0.10] | 0.87 [0.86, 1] | 1, 420 | 2848.93 | < **0.001** |
| | P:R | 0.98 [0.14] | 0.10 [0.016 1] | 1, 420 | 48.90 | < **0.001** |

#### B. F2 weight and length at juvenile and adult stages

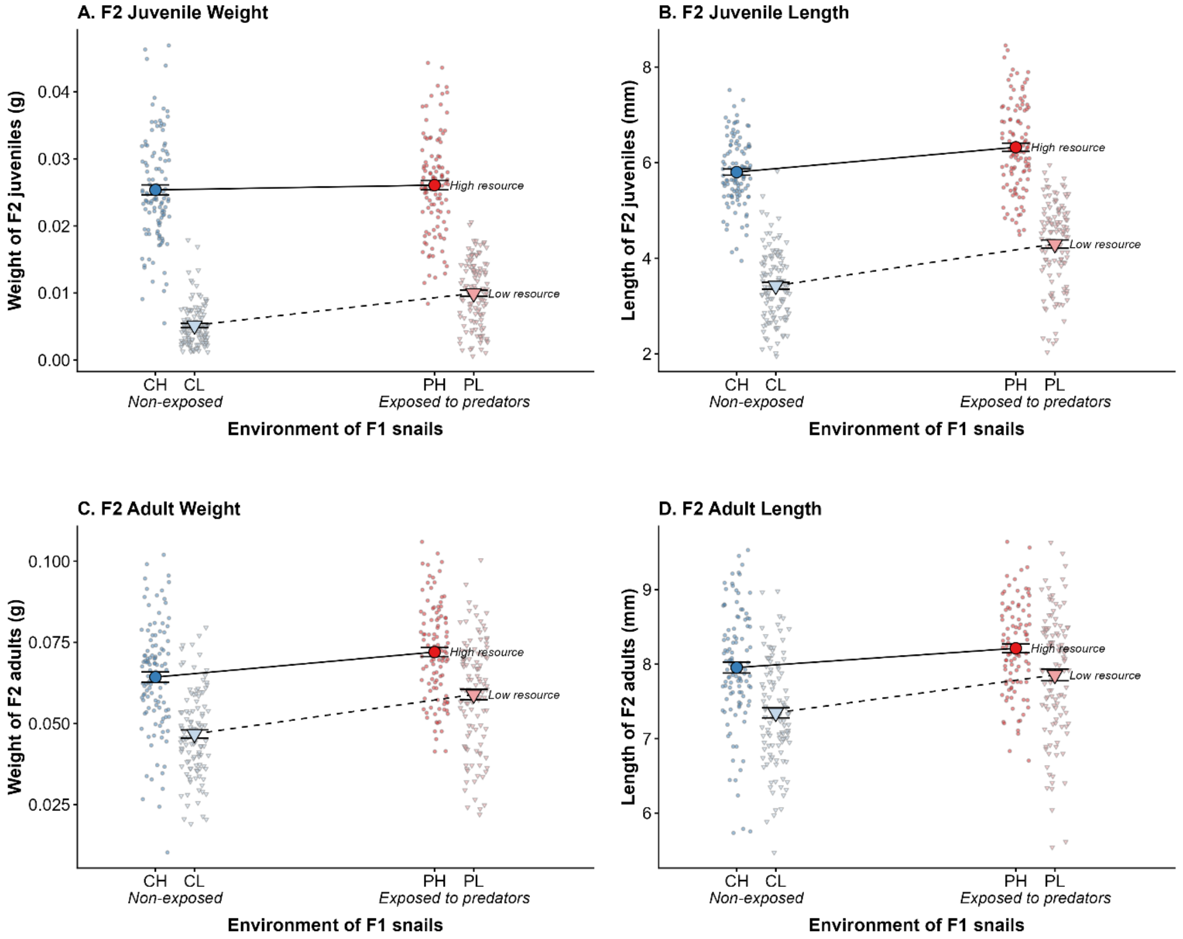


F2 snail weight and length at the juvenile and adult stages were influenced by the interaction between parental predator-cue exposure and parental resource availability (interactions were significant for juvenile length and weight, marginally significant for adult length, and non-significant for adult weight; see Table below). Parental exposure to predator-cues significantly increased the length and weight of F2 snails in most parental resource environments. In the high resource parental environment, juveniles were longer (PH-CH: 9% longer, $t[443] = 4.72$, $p < 0.001$) but had a similar weight ($t[436] = 0.85$, $p = 0.832$), while adults were longer and heavier (PH-CH: 3% longer, $t[417] = 2.47$, $p = 0.066$; 12% heavier, $t[431] = 3.68$, $p = 0.002$). In the low resource parental environment, both juveniles and adults from exposed parents were longer and heavier (juvenile PL-CL: 35% longer, $t[443] = 7.96$, $p < 0.001$ and 92% heavier, $t[436] = 5.80$, $p < 0.001$; adult PL-CL: 7% longer, $t[417] = 5.00$, $p < 0.001$ and 26% heavier, $t[431] = 5.81$, $p < 0.001$). Overall, F2 juvenile and adult snails from the low resource parental environment were significantly shorter and lighter than F2 ones from the high resource parental environment (juvenile CL-CH: 41% shorter, t[443] = -21.62, p < 0.001 and 80% lighter, t[436] = -24.42, p < 0.001; juvenile PL-PH: 32% shorter, t[443] = -18.44, p < 0.001 and 62% lighter, t[436] = -19.30, p < 0.001; adult CL-CH: 8% shorter, t[417] = -5.98, p < 0.001 and 27% lighter, t[431] = -8.39, p < 0.001; adult PL-PH: 4% shorter, t[417] = -3.42, p = 0.004 and 18% lighter, t[431] = -6.22, p < 0.001).

| Trait | Effect | Estimate [SE] | Effect size $\eta^2$[CI] | NumDf , DenDf | F_value | P_value |
|---|---|---|---|---|---|---|
| **F2 Juvenile weight** | Predation | 0.001 [0.001] | 0.04 [0.02, 1] | 1, 436 | 22.31 | **< 0.001** |
| | Resource | -0.020 [0.001] | 0.69 [0.65, 1] | 1, 436 | 956.66 | **< 0.001** |
| | P:R | 0.004 [0.001] | 0.03 [0.01, 1] | 1, 436 | 12.03 | **< 0.001** |
| **F2 Juvenile length** | Predation | 0.52 [0.11] | 0.16 [0.11, 1] | 1, 443 | 80.37 | **< 0.001** |
| | Resource | -2.38 [0.11] | 0.64 [0.60, 1] | 1, 443 | 802.05 | **< 0.001** |
| | P:R | 0.35 [0.15] | 0.01 [0.00, 1] | 1, 443 | 5.18 | **0.023** |
| **F2 Adult weight** | Predation | 0.008 [0.002] | 0.09 [0.05, 1] | 1, 431 | 45.06 | **< 0.001** |
| | Resource | -0.02 [0.002] | 0.20 [0.15, 1] | 1, 431 | 106.88 | **< 0.001** |
| | P:R | 0.004 [0.003] | 0.00 [0.00, 1] | 1, 431 | 2.24 | 0.135 |
| **Fé Adult length** | Predation | 0.26 [0.10] | 0.06 [0.03, 1] | 1, 417 | 28.18 | **< 0.001** |
| | Resource | -0.61 [0.10] | 0.10 [0.06, 1] | 1, 417 | 44.53 | **< 0.001** |
| | P:R | 0.25 [0.14] | 0.00 [0.00, 1] | 1, 417 | 2.94 | 0.087 |

**Supplementary material 3:** Accounting for mass in reproductive outputs (analyses of covariances)

**F1 generation** - Egg and juvenile numbers produced by the F1 adult snails were independently influenced by resource availability and total mass, with no significant interaction between predator-cue exposure and resource availability when total mass was accounted for (Eggs – Mass: $\chi^2$ (1, N = 419) = 15.81, p < 0.001; Predation: $\chi^2$ (1, N = 419) = 1.08, p = 0.300; Resource: $\chi^2$ (1, N = 419) = 16.33, p < 0.001; Predator x Resource: $\chi^2$ (1, N = 419) = 0.12, p = 0.733; Juveniles - Mass: $\chi^2$ (1, N = 419) = 11.60, p = 0.001; Predation: $\chi^2$ (1, N = 419) = 1.36, p = 0.243; Resource: $\chi^2$ (1, N = 419) = 20.87, p < 0.001; Predator x Resource: $\chi^2$ (1, N = 419) = 0.01, p = 0.929).

**F2 generation** - Egg number produced by the F2 adult snails was not influenced by parental predator-cue exposure or parental resource availability when total mass was accounted for (Mass: $\chi^2$ (1, N = 430) = 60.32, p < 0.001; Predator: $\chi^2$ (1, N = 430) = 0.31, p = 0.578; Resource: $\chi^2$ (1, N = 430) = 0.24, p = 0.627; Predator x Resource: $\chi^2$ (1, N = 430) = 1.38, p = 0.240). Juvenile number produced by the F2 adult snails was significantly influenced by parental resource availability and total mass, with no significant interaction between parental predator-cue exposure and parental resource availability (Mass: $\chi^2$ (1, N = 430) = 31.39, p < 0.001; Predator: $\chi^2$ (1, N = 430) = 0.36, p = 0.546; Resource: $\chi^2$ (1, N = 430) = 51.01, p < 0.001; Predator x Resource: $\chi^2$ (1, N = 430) = 2.14, p = 0.143). F2 snails from the low resource parental environment produced a higher number of juveniles, regardless of parental exposure to predator cues.
